# Long-read transcriptomics of Ostreid herpesvirus 1 uncovers a conserved expression strategy for the capsid maturation module and pinpoints a mechanism for evasion of the ADAR-based antiviral defence

**DOI:** 10.1101/2024.05.03.592320

**Authors:** Umberto Rosani, Enrico Bortoletto, Xiang Zhang, Bo-Wen Huang, Lu-Sheng Xin, Mart Krupovic, Chang-Ming Bai

## Abstract

Ostreid herpesvirus 1 (OsHV-1), a member of the family *Malacoherpesviridae* (order *Herpesvirales*), is a major pathogen of bivalves. However, the molecular details of the malacoherpesvirus infection cycle and its overall similarity to the replication of mammalian herpesviruses (family *Orthoherpesviridae*) remain obscure. Here, to gain insights into the OsHV-1 biology, we performed long read sequencing of infected blood clams, *Anadara broughtonii*, which yielded over one million OsHV-1 long reads. This data enabled the annotation of the viral genome with 78 gene units and 274 transcripts, of which 67 were polycistronic mRNAs, 35 ncRNAs and 20 natural antisense transcripts (NATs). Transcriptomics and proteomics data indicate preferential transcription and independent translation of the capsid scaffold protein as an OsHV-1 capsid maturation protease isoform. The conservation of this transcriptional architecture across *Herpesvirales* likely indicates its functional importance and ancient origin. Moreover, we traced RNA editing events using short read sequencing and supported the presence of inosine nucleotides in native OsHV-1 RNA, consistent with the activity of ADAR1. Our data suggests that, whereas RNA hyper-editing is concentrated in specific regions of the OsHV-1 genome, single nucleotide editing is more dispersed along OsHV-1 transcripts. In conclusion, we revealed the existence of a conserved pan-*Herpesvirales* transcriptomic architecture of the capsid maturation module and uncovered a transcription-based viral counter defence mechanism presumably facilitating the evasion of the host ADAR antiviral system.

**Author Summary:** Ostreid herpesvirus 1 (OsHV-1, family *Malacoherpesviridae*) is a major pathogen of bivalve species, causing devasting mortalities and substantial economic losses of aquaculture species. The divergence of OsHV-1 compared to more extensively studied mammalian herpesviruses (family *Orthoherpesviridae*) hampered the understanding of its biology. We performed a deep characterization of the OsHV-1 transcriptome based on long-read RNA sequencing produced from experimentally infected blood clams (*Anadara broughtonii*). Owing to the superior power of long read sequencing to disentangle overlapping transcript isoforms, we could reveal the complexity of the OsHV-1 transcriptome, composed of 274 transcripts. Despite the extensive divergence of OsHV-1 from vertebrate herpesviruses, we reported the presence of a pan-*Herpesvirales* transcriptomic architecture of the capsid maturation module, likely underpinning a conserved functional role in capsid assembly. Furthermore, we revealed the peculiar OsHV-1 transcriptomic patterns, presumably facilitating the evasion of the ADAR anti-viral defence system. In particular, OsHV-1 generates “molecular decoys” by co-expressing sense-antisense transcripts that sequester most ADAR RNA hyper-editing. Both these aspects support the existence of a functional role of “transcriptional architecture” in OsHV-1, contributing to a better understanding of the molecular behaviour of this virus.

## Introduction

Ostreid herpesvirus 1 (OsHV-1, family *Malacoherpesviridae*) is one of the two described herpesviruses infecting invertebrates, including scallop, clam and oyster species (1). OsHV-1 is a major pathogen of bivalves, threatening aquaculture production due to its environmental persistence and multi-host infection (2). Massive mortalities of Arc clam *Anadara broughtonii* (previously known as *Scapharca broughtonii*) associated with OsHV-1 have been reported in China since 2012 (3), whereas the OsHV-1 micro-variant has been linked to mortalities in spat and juvenile *Crassostrea gigas* reported in France since 2008, with devasting economic and social impacts (2,4). The recent report of malacoherpes-like viruses circulating in coastal waters poses concerns for possible future zoonoses and calls for better understanding of virus-host interactions in the megadiverse group of molluscs (5). Thus, a better understanding of the molecular details of the virus replication cycles, including the interplay between antiviral defence and counter defence mechanisms (6,7), is of utmost importance. One way to achieve this goal is by integrating different state-of-the-art ‘omics’ approaches (8,9), as demonstrated for SARS-CoV-2 (10), human immunodeficiency virus type 1 (11), and Crimean–Congo haemorrhagic fever virus (9).

The OsHV-1 infection process has been investigated in *C. gigas* and *A. broughtonii* using classical and high-throughput sequencing approaches (12–17). Soon after the first mortality episode reported in France, the complete OsHV-1 genome was sequenced and used to substantiate its taxonomic classification among herpesviruses; however, the high divergence from all known members of the order *Herpesvirales* suggested that it belongs to a new family, subsequently named *Malacoherpesviridae*. The family was expanded to include a second member, Haliotid herpesvirus 1 (HaHV-1), which infects gastropod species (1,18). The evolutionary history of *Malacoherpesviridae* and their relationship to vertebrate herpesviruses remain obscure due to high sequence divergence and paucity of mechanistic studies on their infection cycle (19,20). A low-coverage PacBio transcriptome map of OsHV-1 and HaHV-1 revealed an unexpected complexity of viral gene arrays (21), suggesting that part of the viral transcription program has been previously overlooked. Arguably, advances in long-read RNA sequencing are now providing the technological platform to resolve complex gene arrays, typical of the densely-coding genomes of some viruses (22). Additionally, the possibility to sequence native RNA directly (Direct RNA Sequencing, DRS) opens new avenues for epitranscriptomic analysis in virology (23,24).

Aiming to gain insights into the OsHV-1 biology, we constructed six native RNA libraries and sequenced them at high-coverage to characterize the OsHV-1 transcriptional program. The long-read data was coupled with short-read and proteomic data to (i) reveal the complexity of the OsHV-1 transcriptional program, including alternative transcription start and stop sites, read-through transcription events and natural-antisense transcripts (NATs), (ii) identify transcriptional architecture with possible functional roles for viral replication and/or interaction with the host, and (iii) validate the extent and distribution of RNA editing events mediated by the host Adenosine Deaminase acting on dsRNA (ADAR), an enzyme deaminating adenosine to inosine and restricting the replication of diverse RNA and DNA viruses, including mammalian herpesviruses (25).

## Results

An OsHV-1 homogenate was injected in a batch of wild *A. broughtonii* specimens, and haemocyte samples were collected from three animals at 6, 12, 24, 36, 48, 60, and 72 hours post-infection (hpi) to extract RNA and proteins. Short-read data (described in (21)) showed that OsHV-1 RNA reads are barely detectable until 24-36 hpi, slightly increase in number at 48 hpi, and become prominent, occupying a considerable part of the sequenced transcriptome at 60 and 72 hpi before the appearance of mortality signs (Figure S1). Accordingly, to ensure the abundance of viral reads in the DRS data, we selected three biological replicates at 60 and 72 hpi and produced more than 30 million (M) of long reads in total (Table 1). The majority of them (92.6-95.6%) are ‘on-target’, i.e., could be mapped to either the Ark clam or OsHV-1 genomes, whereas ‘off-target’ reads were possibly produced from other biotic components, sequencing artifacts or from the transcription of host genes not present in the reference genome (Figure 1a). DRS data included fewer off-target reads compared to the paired Illumina datasets, possibly because of the enrichment for polyadenylated transcripts, a procedure not applied for Illumina total RNA sequencing. An average of 2.2 and 4.2% of the DRS reads at 60 and 72 hpi, respectively, mapped to the OsHV-1 genome (Figure 1b), showing a good correlation with Illumina data (R^2^ =0.83, Figure 1c). Mass-spectrometry (MS) analysis performed on protein extracts collected throughout the course of infection generated a total of 798,086 spectra, of which 0.09% (147 unique peptides) mapped to the predicted OsHV-1 proteome.

**Figure 1.**
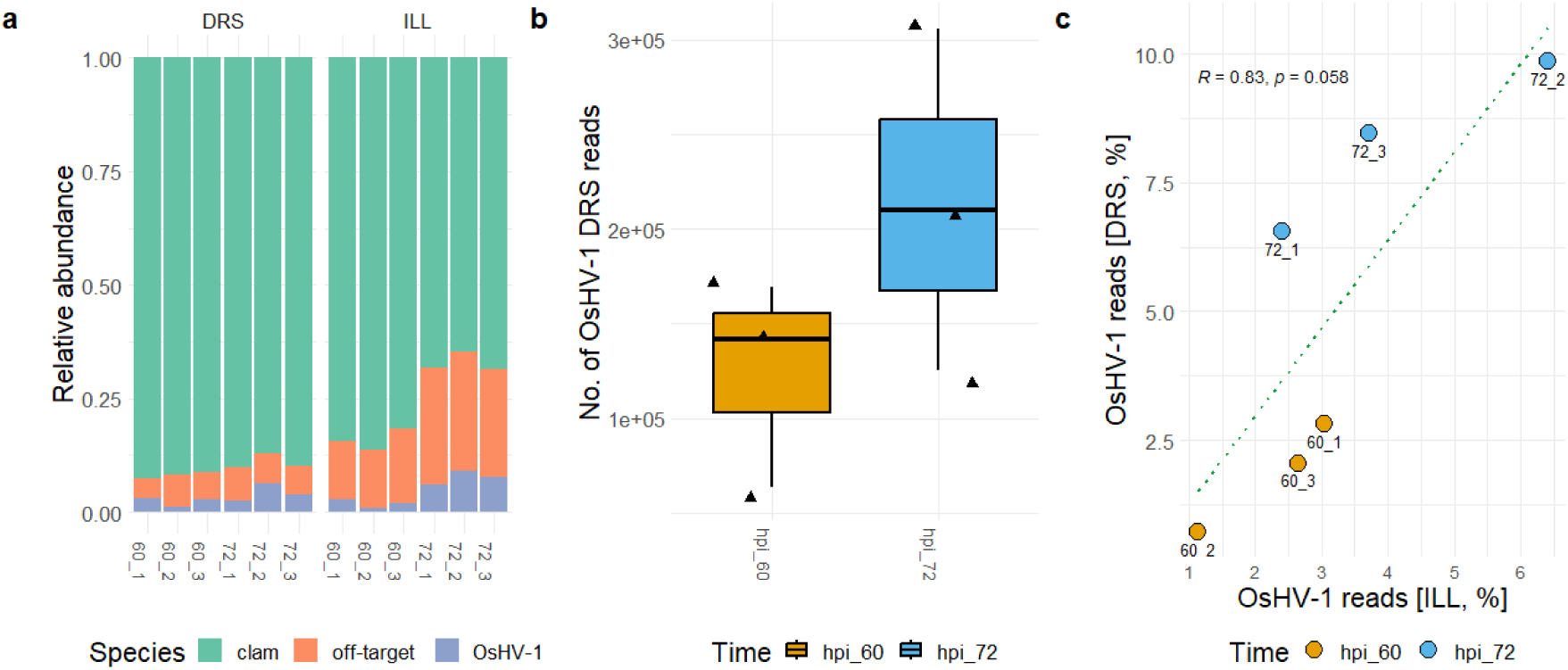
Sequencing of the OsHV-1 transcriptome using different platforms. (a) Relative abundance of the on-target and off-target reads in the six DRS and Illumina (ILL) datasets. The fractions of reads mapping on the *A. broughtonii* genome are reported in green, OsHV-1 in purple and off-target reads in orange. (b) Number of DRS reads mapped on the OsHV-1 genome at 60 and 72 hpi. (c) Distribution of the fraction of Illumina and DRS reads mapped on the OsHV-1 genome in the six sample. The Spearman correlation value and associated p-value are reported on the graph.

**Table 1.**
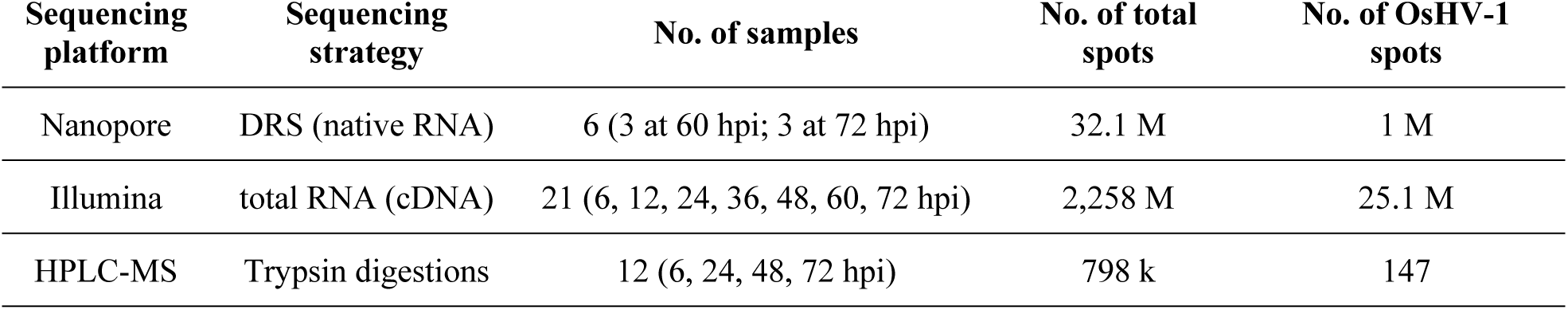
Data statistics.

| Sequencing platform | Sequencing strategy | No. of samples | No. of total spots | No. of OsHV-1 spots |
| --- | --- | --- | --- | --- |
| Nanopore | DRS (native RNA) | 6 (3 at 60 hpi; 3 at 72 hpi) | 32.1 M | 1 M |
| Illumina | total RNA (cDNA) | 21 (6, 12, 24, 36, 48, 60, 72 hpi) | 2,258 M | 25.1 M |
| HPLC-MS | Trypsin digestions | 12 (6, 24, 48, 72 hpi) | 798 k | 147 |

### Long-read transcriptomics of the OsHV-1 infection

The sequenced transcriptome of OsHV-1 included a total of 1,015,000 DRS reads. The detection of sudden coverage changes (increase or decrease) on each strand of the viral genome was used to identify 292 Transcription Start Sites (TSS) and 201 Transcription Termination Sites (TTS) (Figure 2a). These TSSs and TTSs were in silico combined per strand to obtain 2,274 pseudo-transcripts, with a length range of 250-15,000 nt. Pseudo-transcripts were used as mapping references for the DRS data and 274 of them were supported by at least three full-length DRS reads and were considered as bona fide transcripts (Figure 2a).

**Figure 2.**
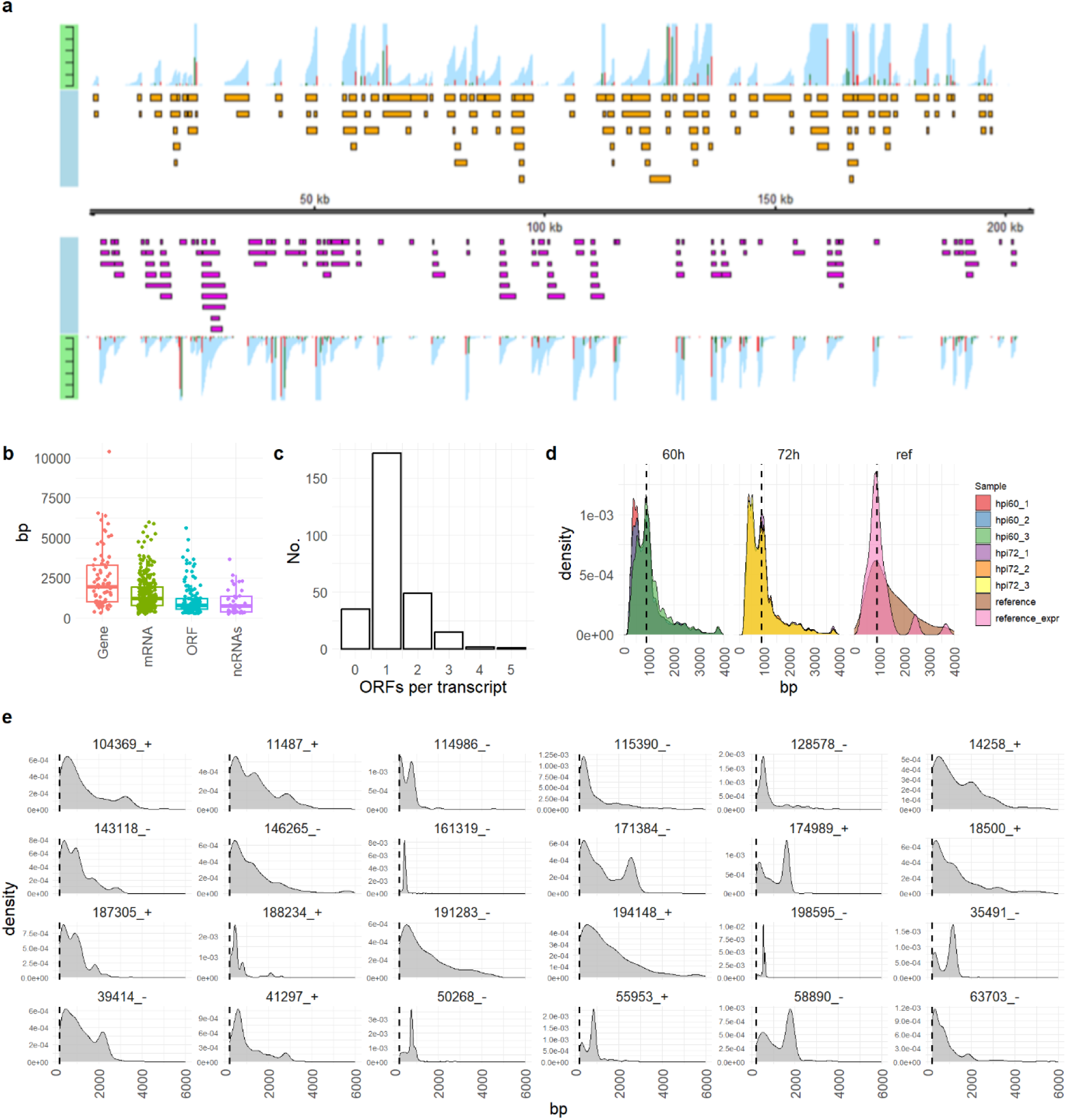
The OsHV-1 transcription map. (a) The identified TSSs (green bars) and TTSs (red bars) are depicted on each strand of the OsHV-1 genome, together with the 274 transcripts reported by orange (forward strand) and magenta (reverse strand) boxes. The height of the TSS and TTS bars represented the frequency of the events and the light-blue overlay represented the coverage of each transcript based on the DRS data. (b) Length distributions of the annotated mRNAs, ORFs, ncRNAs and gene units of OsHV-1. (c) Distribution of the number of predicted ORFs per transcript. Hits with no ORF referred to ncRNAs. (d) Distribution of the sizes of the OsHV-1 DRS reads at 60 and 72 hpi, reported together with the sizes of all the 276 predicted transcripts (“reference” in legend) and of the most expressed ones (“reference_expr”). A dotted line denotes the size of the main peak (976 bp). (e) Density plot referring to the lengths of the DRS reads mapped to 8 selected genes among the 25 showing RTT evidence. The length range reported in the graph started from 200 nt after the TTS. The panel is a selection of the results reported in Figure S2.

Thirty-five of them did not contain ORFs and were classified as non-coding RNAs (ncRNAs), with a size ranging from 0.26 to 3.6 kb (Figure 2b-c). Additionally, 67 transcripts were identified as polycistronic, encoding between two and five ORFs each. By collapsing overlapping transcripts for each genome strand, 78 gene units were defined, characterized by longer sizes compared to transcripts or coding regions (Figure 2b). A total of 328 proteins were predicted to be encoded by the 274 OsHV-1 transcripts, reduced to 133 after removing identical hits originating from gene isoforms.

Our analysis confirmed the expression of 113 out of the 131 ORFs previously annotated in the reference OsHV-1 genome (Table S1). The genes for which transcripts were not obtained include ORF1 and ORF2, located within the repeated terminal region of the genome and therefore present in two copies, and ORF5, ORF16, ORF18, ORF27, ORF36, ORF69, ORF74, ORF81, ORF105, ORF116, ORF118, ORF119, ORF120 and ORF121, all encoding functionally uncharacterized proteins. Notably, 13 ORFs located on 9 transcripts are not annotated in the reference genome, although some of these were annotated in non-reference OsHV-1 strains (e.g. ORF60_x reported in OsHV-1-PT), whereas ORFs encoded antisense with regard to the reference annotations were missed previously altogether.

The size distribution of the OsHV-1 DRS reads is reproducible among five of the six samples (Figure 2d), with all the samples showing a bimodal size distribution. The first peak matched the average length of the most expressed transcripts (Figure 2d), whereas the second peak, corresponding to shorter transcripts, likely represented RNA degradation products or incomplete transcription events. Notably, only sample 60-hpi_3 possessed a higher “full-length peak” compared to the short one. In order to verify this observation, we mapped the DRS reads to the 78 gene units separately and evaluated the distribution of the read lengths compared to the nominal length of each gene. As expected, for most of the genes, the DRS size distribution discretely peaked near the annotation boundaries or, occasionally, before them if shorter gene isoforms are expressed (Figure S2). However, for 25 genes, we obtained evidence of Read-Trough-Transcription (RTT), with fractions of the reads exceeding gene boundaries. RTT generally produced molecules of variable lengths, except for a few cases in which we detected peaks of discrete size, possibly corresponding to overlooked isoforms (e.g. gene 174989_+ in Figure 2e).

Next, we investigated the dynamic expression of the OsHV-1 transcriptome based on DRS (60-72 hpi) and short read data (6-72 hpi). DRS data revealed that late in infection the most expressed transcripts are generally conserved in 5 samples, whereas sample 60_3 showed a higher number of transcripts with a minimal expression level of 10,000 Transcripts Per Millions (TPMs), resulting in a different global expression profile (Figure 3a). Two of the top5 expressed transcripts per sample are present in all six datasets (127600_128424, a gene encoding the putative capsid protein VP23 and 64787_65654 with unknown function), two in five datasets (23934_24291 and 166004_166879, the first being a ncRNA and the second encoding a short isoform of the capsid maturation protease), one in four datasets (135399_136225, unknown function) and four transcripts are in the top5 list of a single sample only. Comparison of the six short read datasets paired to DRS data showed that the correlation of expression levels between the two technologies is generally good, in particular for the transcripts with mid-to-high expression levels (Figure S3). For 48 transcripts short read data did not support the expression observed with DRS data, the latter being at least 10 folds higher, likely due to the intrinsic limitation of short reads when dealing with overlapping genes.

**Figure 3.**
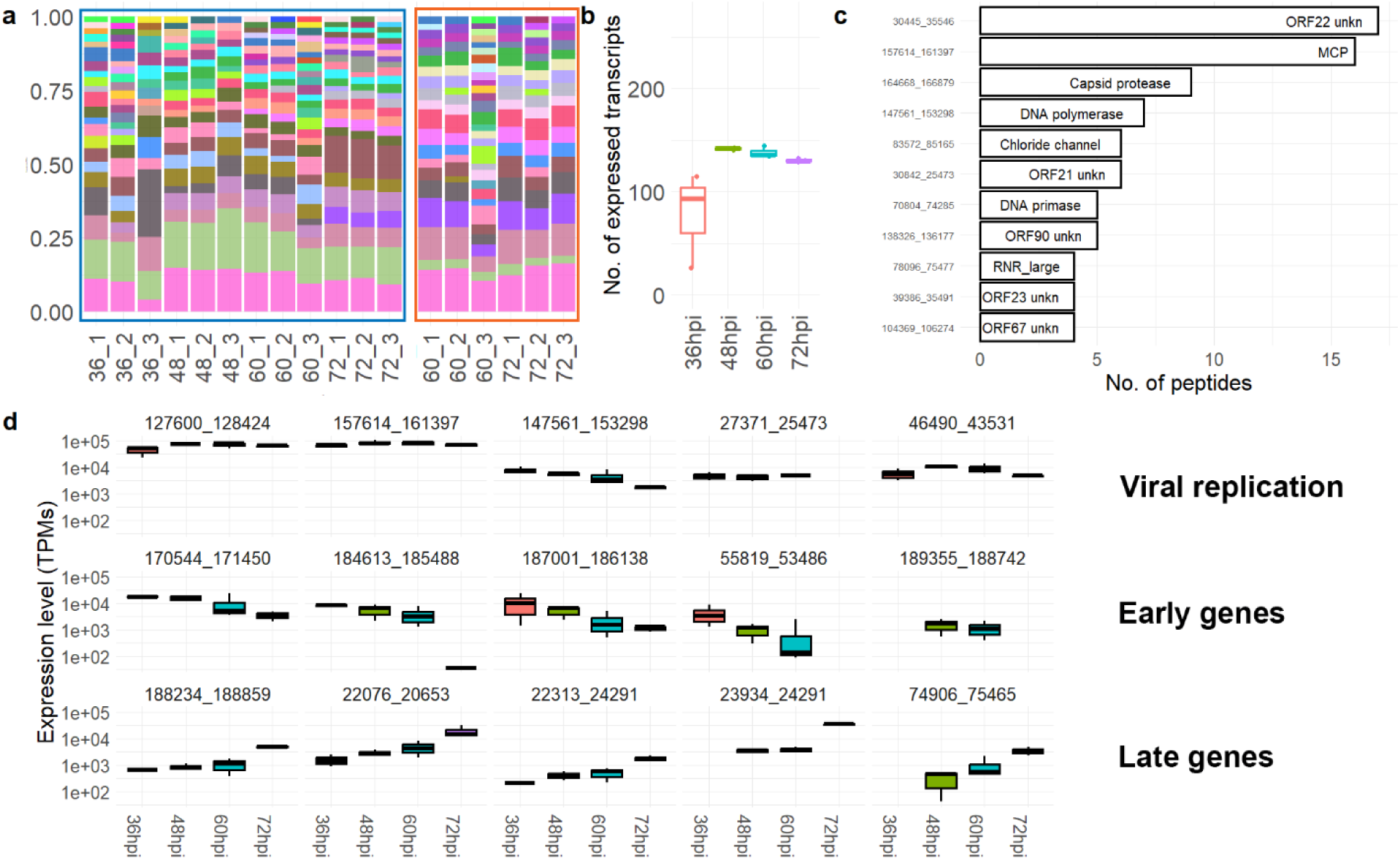
Expression dynamic of OsHV-1. (a) The relative abundances of the OsHV-1 transcripts with expression levels higher than 10,000 Transcripts Per Millions (TPMs) are depicted. The part squared in orange referred to DRS data (60 and 72 hpi), whereas the part squared in blue referred to short read data (36 – 72 hpi). The identity of the samples is indicated in the names of the bars. The expression dataset is included in Table S2. (b) Number of expressed transcripts (> 100 TPMs) in the samples, measured using short reads. (c) The OsHV-1 proteins with at least four matching peptides are reported, with the protein identities indicated into the bars. (d) Expression trends of selected OsHV-1 transcripts referring to viral genome replication and virion morphogenesis genes (VP23, MCP, DNA polymerase, ribonucleotide-reductase small subunit and the portal protein [first row]), early genes (second row) and late genes (third row) are reported averaged per time point.

Extending the expression analysis to earlier time points (6, 12, 24, 36 and 48 hpi) allowed detection of OsHV-1 transcripts starting from 36 hpi (Figure 3b). The expression heatmap clustered the samples into three groups: the first included the 36-hpi samples with a limited number of expressed transcripts compared to later time points; the second included 48 and 60 hpi samples, and the third cluster accommodated the 72 hpi samples. The second cluster is further organized into two subclusters: one for two 48 hpi and 60-hpi_3 samples and the other subcluster included the rest of the samples (Figure S4). The expression of gene encoding major capsid protein (MCP), capsid protein VP23, scaffold protease, portal protein (identified herein as product of ORF28, transcript 46490_43531; Table S1), DNA polymerase, ribonucleotide reductase subunits and several proteins of unknown function were detected early in the infection (Figure 3d and Table S2). During the later stages of infection, the identification of the viral genes became more challenging because of the incremental number of expressed transcripts, possibly linked to the presence of RNAs destined for degradation rather than active translation. Overall, we identified seven putative early transcripts and five late ones (Figure 3d). Notably, none of these stage-specific transcripts encoded for proteins with a predictable function. The 147 peptides matching the OsHV-1 genome found by MS proteomics originated from 41 proteins. The top three most abundant detected proteins were ORF22 (unknown function), MCP (ORF104) and capsid maturation protease (ORF107) (Figure 3c).

### A transcriptional architecture of the capsid maturation protease is conserved among distant herpesviruses

Virion formation is a quintessential feature, which distinguishes viruses from cellular organisms and all other types of mobile genetic elements (26). Given the importance of virion morphogenesis genes and our observation that they are among the most strongly expressed, we investigated in detail the OsHV-1 regions coding for MCP (ORF104), capsid proteins VP19 (ORF112) and VP23 (ORF82), portal protein (ORF28) and capsid maturation protease (ORF107). The functions of these proteins have not been experimentally validated, thus similarity searches were used to support their identities (File S1).

The genomic region coding for the MCP included two isoforms, with two different TSSs and one shared TTS, resulting in either the full-length MCP (MCP_1) or a 5’-truncated MCP (MCP_2) (Figure 4a), lacking the first 471 amino acids. Although MCP_1 and MCP_2 showed similar expression levels in the samples (Figure 4f), the TSS of MCP_1 showed a higher frequency and proteomic data supported the presence of the full-length protein, encoding both the floor and the tower domains. The putative capsid protein VP19, which we identify as a homolog of orthoherpesviral triplex 1 protein, is encoded as the first ORF of a bicistronic transcript, characterized by four isoforms coding either for both ORFs (VP19 and ORF113; referred to as VP19_polycist) or for VP19 (1 isoform) and ORF113 (2 isoforms) separately (Figure 4b). The ORF113 isoforms and VP19_polycist appeared highly expressed compared to the standalone VP19 isoform (Figure 4f). The genomic region coding for the capsid protein VP23, a homolog of the orthoherpesviral triplex 2 protein, showed a single isoform with the highest expression levels in all samples, greatly exceeding the other considered genes (Figure 4c). The gene coding for the portal protein included two isoforms, a full-length protein of 853 amino acid residues (named “portal” in Figure 4d) and an isoform coding for a shorter in-frame ORF, lacking the first 600 residues. Notably, the shorter isoform showed an average four-fold higher expression than the full-length isoform (Figure 4f). The capsid maturation protease gene displays an even higher transcriptional complexity, including eight overlapping transcripts. The transcript encoding the full-length protein (164668_166879, 689 aa, labelled “Capsid protease” in Figure 4e, light-blue arrow) embedded a shorter in-frame ORF, potentially encoding an N-terminally truncated protein of 367 aa residues (named “Scaffold” in Figure 4e, green arrow). Both transcripts also showed 3’-RTT, sharing the TTS with a distal gene, also characterized by two isoforms (named “Capsid protease polycist” and “Scaffold polycist”, and distal 1 and 2, respectively in Figure 4e). Finally, two ncRNAs are present (purple arrows in Figure 4e). Gene expression levels based on DRS data indicated that ncRNA_2 and the scaffold transcript are the most expressed ones (Figure 4f), as also evident from the high frequencies of their TSSs and shared TTS (Figure 4e). The capsid protease showed more limited expression levels, as well as the polycistronic transcripts. Notably, all the considered isoforms involved in capsid assembly and maturation showed a limited variation in their relative expression between samples, possibly indicating that they function in stoichiometric quantities. These results suggest that tight regulation of the capsid assembly and maturation genes is ensured by several different mechanisms.

**Figure 4.**
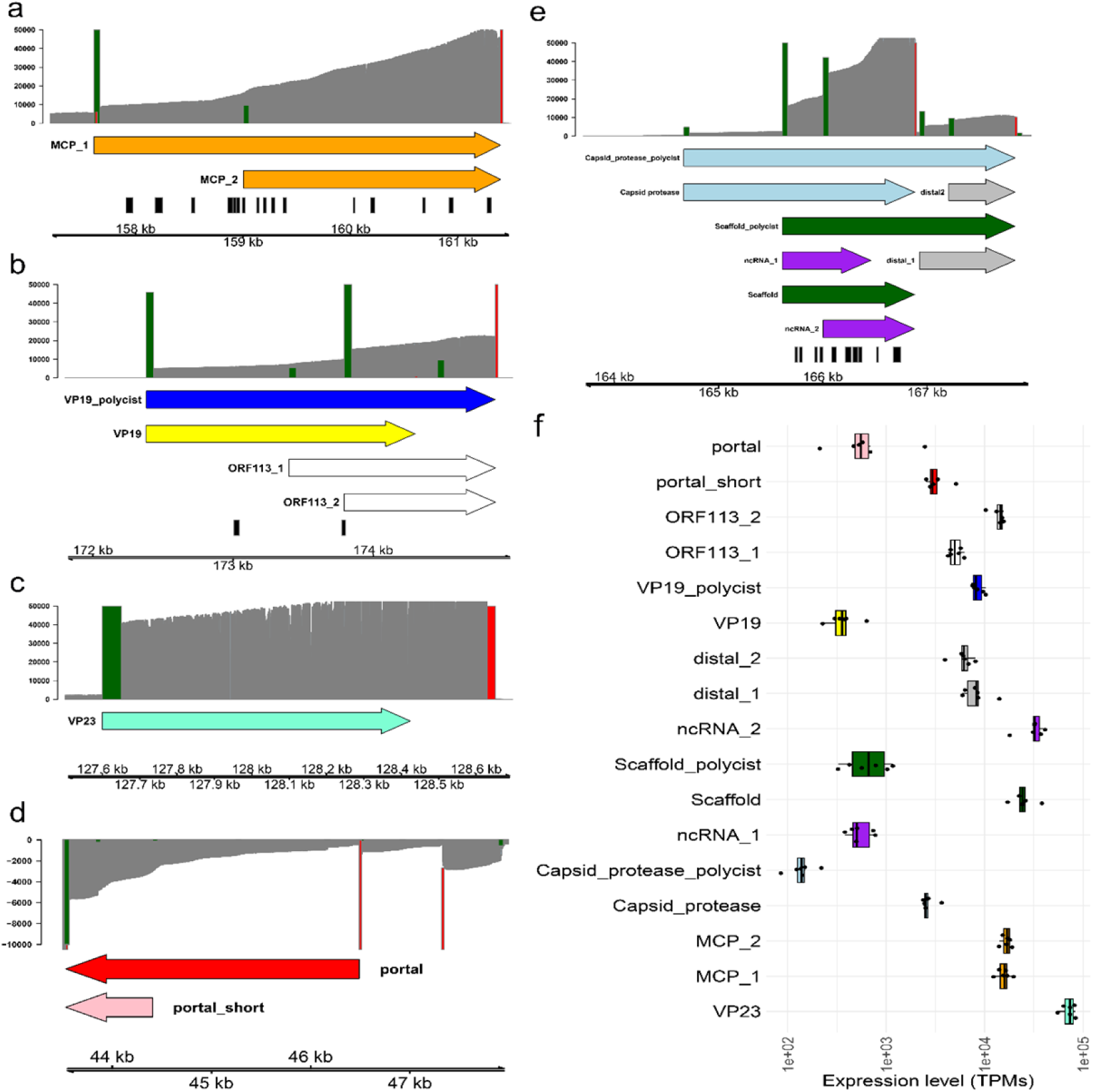
OsHV-1 capsid assembly and maturation. (a) The transcriptional landscape of the OsHV-1 major capsid protein (MCP) is reported, with arrows depicting the isoforms, together with their TSSs and shared TTS (green and red bars, respectively). The DRS coverage is reported as grey background, and the peptide positions identified by proteome analysis are depicted by black bars. (b) Transcriptional landscape of VP19 gene region. (c) Transcriptional landscape of VP23 gene region. (d) Transcriptional landscape of the portal protein. (e) Transcriptional landscape of the capsid protease of OsHV-1 (164,000-168,000, forward strand). (f) Expression levels were reported as TPMs for the considered isoforms in the six samples. The colour code in (f) matched the colours used in the other panels.

### ADAR-mediated editing and hyper-editing along OsHV-1 genome

The genomic positions of the 274 predicted OsHV-1 transcripts, collapsed into the 78 gene units, were mined to identify 20 Natural Antisense Transcripts (NATs) and 10 regions of overlapping sense-antisense transcription. Regions 4, 5, 6, 7, 8 and 9 are characterized by divergent overlaps, i.e. overlaps of the 5’-ends of the transcripts, regions 1 and 2 contained antisense embedded genes and regions 3 and 10 corresponded to convergent overlaps, i.e. overlaps of transcripts’ 3’-ends (Figure 5a). Detailed characterization of the expression levels of these NATs revealed that 9 out of 10 NAT pairs showed medium-to-high expression levels in the six samples, with three pairs exceeding an average of 10,000 TPMs (Figure 5b).

**Figure 5.**
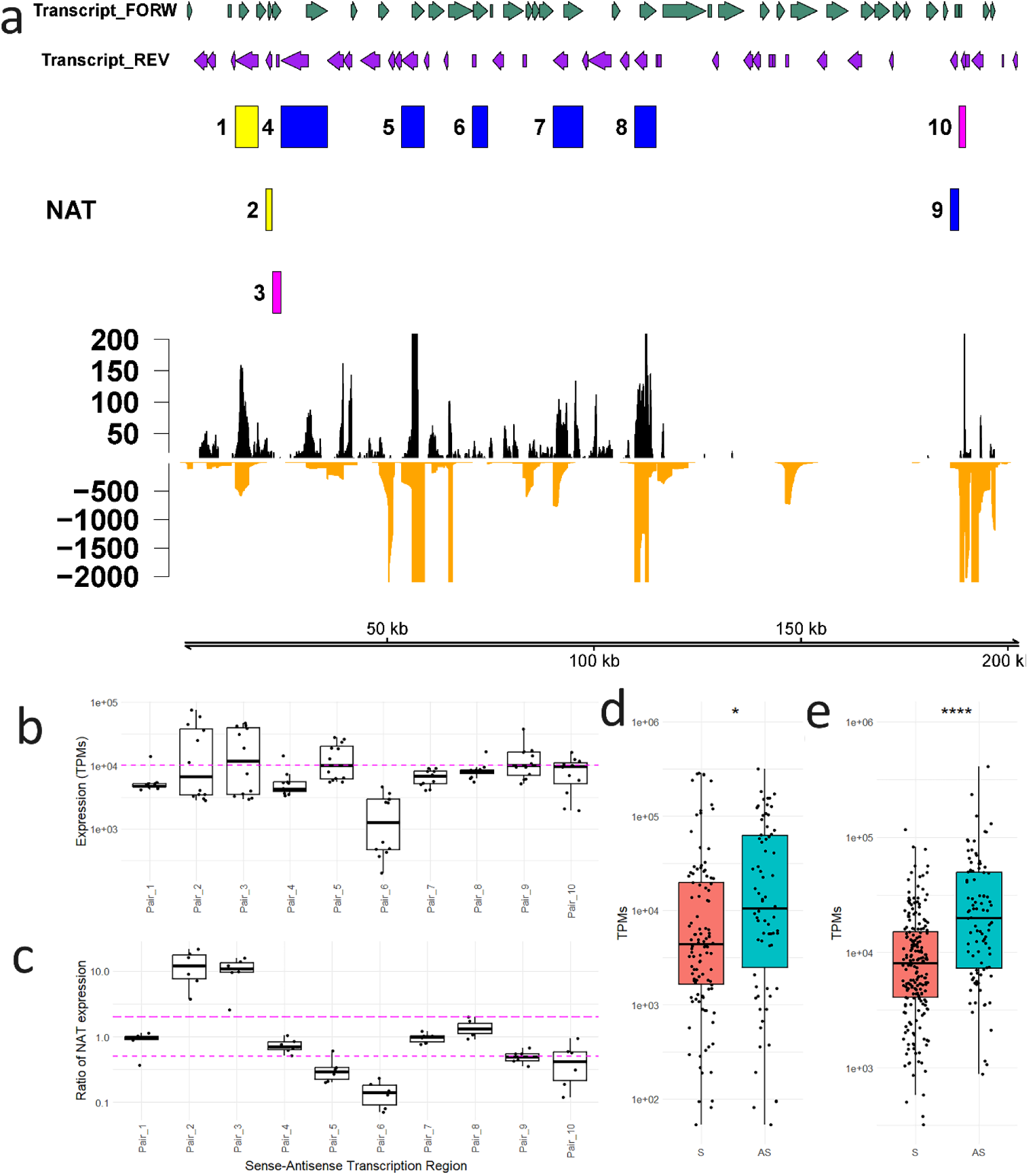
Antisense transcription along the OsHV-1 genome and RNA editing. (a) The figure depicted the transcriptional landscape of OsHV-1 with forward (green arrows) and reverse (purple arrows) transcripts, the 10 sense-antisense transcription regions with a colour code indicating if they are referring to convergent (pink boxes), divergent (blue boxes) or embedded (yellow boxes) NAT overlaps as well as the distribution of hyper-edited short reads (black profile) and of edited DRS reads (orange profile). (b) Expression levels of the NATs measured as TPMs and divided by the 10 sense-antisense transcription pairs. The magenta dotted line indicated the 10,000 TPMs level (c) The expression ratio between the sense and the antisense NATs for each pair is reported. The magenta lines indicated the range of balanced expression between NATs (5 folds). (d) Distribution of the number of edited DRS reads between sense transcripts (S) and NATs (SA). *: p-value < 0.05, t-test. (e) Distribution of the number of hyper-edited reads between sense transcripts (S) and NATs (SA). ****: p-value < 0.001, t-test.

To gain insights into the potential roles of the NATs, we considered the distribution of polymorphisms across different OsHV-1 transcripts, which could signify editing by host defence systems, and of the extent of ADAR-mediated adenosine-to-inosine modifications in OsHV-1. This was obtained by analysing the distribution of inosines and of hyper-edited reads. The mapping of short read data on the OsHV-1 genome was also used to identify 465 ADAR-compatible single nucleotide variations (SNVs, A-to-G or T-to-C), located predominantly in non-coding regions (65.9%), whereas 20.2% caused non-synonymous changes and 13.9% synonymous changes (Table S3). A total of 226 SNVs were validated for the presence of inosine, using at least five DRS reads with a minimal strandedness of 2 folds (i.e., A-to-G variations supported by DRS reads mapping in forward and T-to-C variations with DRS reads mapping in reverse). Overall, 60,420 RNA molecules sequenced by DRS contained at least one inosine (5.9% of total reads), with a slight reduction at 72 hpi (Figure S5b). Notably, sample 72_1 comes across as an outlier (sample 72_1), with a higher hyper-editing level and a lower editing level, in comparison with the paired samples. In parallel, we extracted from the six Illumina datasets 21,180 hyper-edited reads, accounting for 0.1% of the total reads, with an increase observed from 60 to 72 hpi (Figure S5a). The distribution of hyper-edited reads and of DRS reads containing inosines was comparable and mostly matched sense-antisense transcription regions 1, 5, 8, 9 and 10 (Figure 5a). An additional hotspot of editing is found around position 150 kb, but this is not supported by hyper-edited reads.

Notably, seven pair of the described NATs, which showed balanced expression between sense and antisense transcripts (Figure 5c), were the most impacted by editing (regions 1, 4, 5, 7, 8, 9 and 10). We also found that hyper-editing preferentially impacted sense-antisense transcription regions, whereas this preference was less pronounced for the single-nucleotide editing traced by the DRS data (Figure 5d-e).

Focusing on the edited DRS reads, most of them possessed a single inosine (52%), with a minority having multiple inosines, up to 36, with no difference among samples (Figure 6a). The average inosine frequency at the edited positions was 10%, with the exception of sample 60-hpi_2 which showed a mean frequency of 25% (Figure 6b). Based on the distribution of inosines along the reads, only 733 DRS reads (1.2%) showed a hyper-editing profile with at least 5 inosines within 150 nt, all located within antisense regions 9 and 10. Eleven transcripts have at least 100 modified reads, but notably only one of them (55953_58943) is a NAT, while most of the others are located near antisense transcription regions (Figure 6c). None of these 11 transcripts encoded proteins with predictable functions. Finally, no significant size differences were found between edited and non-edited reads (Figure 6d).

**Figure 6.**
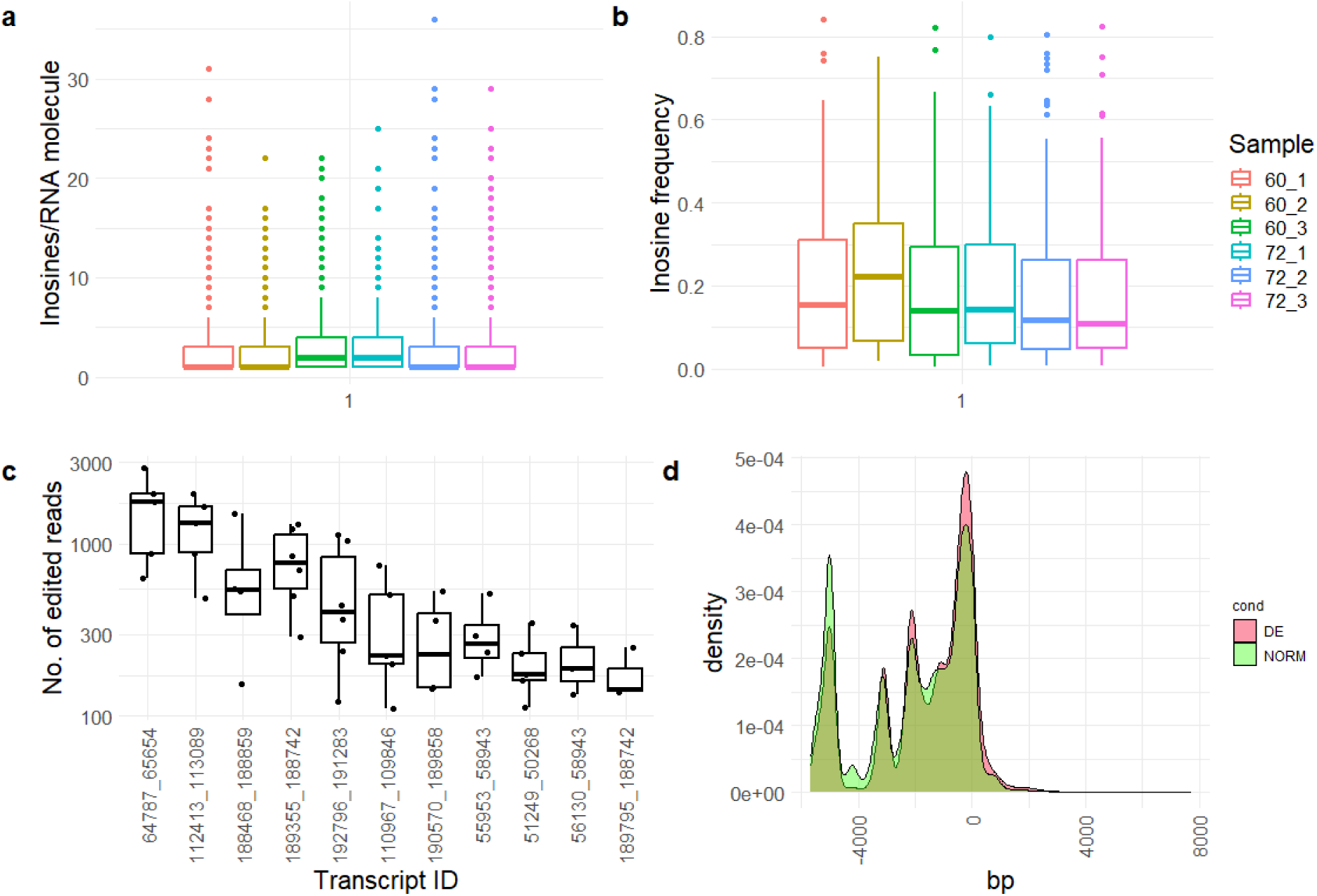
ADAR editing and hyper-editing in OsHV-1. (a) Distribution of the number of inosines nucleotides per RNA molecule in the six samples. (b) Distribution of the frequencies of inosines traced in the 226 validated SNVs measured in the six samples. (c) Number of edited DRS reads per transcripts. Only the 11 transcripts with more than 100 edited reads in at least three samples are reported. (d) Size profiles of the DRS reads (green, NORM) and of the edited DRS reads (red, DE), normalized by the nominal lengths of the matching genes.

In conclusion, the presence of both RNA editing and hyper-editing support the active role of ADAR during OsHV-1 infection. However, hyper-editing and editing seem to follow different trajectories, with the former occurring within regions of perfect sense-antisense matches only, and the latter being more diffuse across the OsHV-1 transcripts.

## Discussion

We performed a deep characterization of the OsHV-1 transcriptome by long read transcriptomics (DRS), complemented with short-read RNA sequencing and MS proteomics. DRS data support and extend by 2.5 folds the number of transcripts identified in the previous OsHV-1 transcriptome map based on PacBio long reads (21). Moreover, our data illuminates interesting new aspects of the OsHV-1 transcription beyond the mere expression analysis. Indeed, we confirm the complexity and high coding density of the OsHV-1 genome, with 274 transcripts grouped into 78 gene units, with evidence of polycistronic transcripts and multiple isoforms involving most of the genes. The high transcriptional complexity has been reported for several representatives of the order *Herpesvirales* (27), making proteomic and/or ribosome profiling data necessary to further investigate the true coding potential of these viruses (28). Our proteomics analysis supported only a small number of OsHV-1 proteins, most of them linked to important viral functions, such as virion formation and genome replication (e.g., the MCP, capsid maturation protease and DNA polymerase) or proteins that, despite having unknown function, were reported as highly expressed in other transcriptomic (13) or proteomic surveys performed in *C. gigas* infected with OsHV-1, such as ORF22 (29).

By leveraging the updated transcriptomic annotation and extending the expression analysis to earlier time points, we captured the temporal dynamics of viral genome expression and categorized viral transcripts into those expressed preferentially in early- or late-infection stages. During the early infection stages, we also detected the expression of genes involved in virion assembly and viral genome replication (e.g., capsid proteins, DNA polymerase). However, the transcripts characteristic of the early or late stages only, were functionally unknown. The considerable fraction of proteins for which functions cannot be predicted highlights the importance of further functional studies which may uncover both new, mollusc-specific aspects of viral biology and mechanisms shared with vertebrate herpesviruses and perhaps even more distantly related non-animal viruses.

Malacoherpesviruses together with vertebrate herpesviruses (families *Orthoherpesviridae* and *Alloherpesviridae*), the recently discovered protist-infecting viruses of the proposed phylum *‘Mirusviricota’* (30) and prokaryotic viruses of the class *Caudoviricetes* (31,32) comprise the realm *Duplodnaviria* (33). Viruses in this vast and ancient realm share the morphogenetic module and form icosahedral capsids from structurally conserved MCPs with the HK97-fold (34). Virion morphogenesis starts with the assembly of an empty capsid which undergoes maturation and is subsequently filled with the viral genomic DNA. A scaffolding protein and the capsid maturation protease, two conserved components of the duplodnavirus morphogenetic module, play key roles in virion morphogenesis (20,35,36). The key proteins for capsid assembly and maturation have been identified by similarity searches in HaHV-1 and OsHV-1 (20). Here, we extended the annotation of the malacoherpesviral morphogenetic module by identifying the OsHV-1 triplex proteins 1 (ORF19) and 2 (ORF23) as well as portal protein (ORF28). The portal is a key component of the virion, which nucleates the capsid assembly and provides a conduit for the entrance of genomic viral DNA (37). The two triplex proteins play an important role in stabilizing the icosahedral capsid and are specific to eukaryotic duplodnaviruses, previously described in vertebrate orthoherpesviruses and protist mirusviruses (30). Notably, similar to other viruses, the two triplex proteins, along with the MCP, are among the most strongly transcribed viral genes (30). Moreover, our DRS data provides evidence that the transcriptional architecture of the scaffolding gene and capsid maturation protease gene is shared between malacoherpesviruses and vertebrate herpesviruses. We found that the capsid maturation protease is encoded as a fusion with the scaffolding protein, and transcription of this gene yields three types of transcripts corresponding to the capsid protease-scaffolding protein fusion and at least one isoform of the standalone scaffolding protein. Notably, the expression levels of the variants are highly different, with the full-length protease-scaffold fusion transcripts showing 10-fold lower levels of expression compared to the scaffolding protein transcript. A similar feature was also reported in vertebrate herpesviruses, HSV-1, bovine alpha-herpesvirus type 1 (BoHV-1) and Marek’s disease virus (MDV) (8,28,38). In BoHV-1, the short isoform of the UL26 gene, called UL26.5, displayed higher expression levels at 12 hours of lytic infection (38), and independent transcription and translation were reported for MDV (28). Our proteome data supported the presence of the capsid scaffold protein, whereas no peptides were detected for the capsid protease, likely due to the low expression of the full-length isoform and autoproteolytic cleavage of the capsid maturation protease. The conservation of the transcriptional architecture and expression profiles in OsHV-1, HaHV-1 (21) and other herpesviruses (8,28,38) suggests its functional importance. Indeed, the covalent fusion with the scaffolding protein would ensure the timely co-incorporation of the protease into empty pro-capsids along with the scaffolding protein. Furthermore, it stands to reason that the amount of the protease necessary for the proteolytic processing of the scaffolding protein would be considerably lower than that of the substrate (i.e., scaffolding protein). Although it is difficult to estimate the divergence time for distantly-related viruses, the pan-*Herpesvirales* conservation of transcriptional architecture of the protease-scaffolding gene module has likely evolved at least in the common ancestor of vertebrate and invertebrate herpesviruses, and hence could predate the divergence of arthropods, molluscs and chordates (39,40).

Our transcriptomic data has also illuminated a new facet of malacoherpesvirus-host interaction involving the ADAR antiviral defence system. We have previously reported ADAR-mediated RNA editing in HaHV-1 and OsHV-1 infecting different host species (21,41), a feature which differentiated malacoherpesviruses from all other known herpesviruses. In fact, in vertebrate herpesviruses, the RNA editing impacts specific genomic features only (e.g. specific miRNAs or transcripts (42)), whereas in malacoherpesvirus, RNA editing extensively impacted multiple transcripts. We previously showed that in OsHV-1, the RNA editing is concentrated within genomic hotspots, whereas in HaHV-1 these modifications are more evenly distributed across the genome (21). However, up to now, the extension of RNA editing was indirectly deduced tracing highly modified short reads, as a proxy of ADAR hyper-editing, whereas no approaches were undertaken to detect inosines. In this study, through the sequencing of native RNA, it became possible to confirm the presence of inosines. So far, only *Dinopore*, which uses a convolutional neural network approach to predict whether a given site is edited (43), and DeepEdit, which, according to the authors, can provide a phased read-level detection of RNA editing events (44), allowed the identification of inosine from DRS data. Despite certain limitations, such as the use of an exogenous model to train the dataset and the requirement of target positions to verify the presence of inosines, based on *DeepEdit*, we could provide independent support for the presence of ADAR editing in OsHV-1 and an accurate evaluation of the extent of such phenomenon.

As result, we confirmed that ADAR hyper-editing mostly occurred within the antisense transcription regions with balanced expression between NATs, because these regions generated abundant dsRNA able to “capture” the enzymatic capability of ADAR proteins, possibly diverting it from functional transcripts, a phenomenon that we previously described as “ADAR decoys” (21). However, here we showed that the distribution of inosines extends beyond NATs, and is mostly associated with single nucleotide editing rather than hyper-editing. This indicates that single-nucleotide editing might be more ‘diffuse’ than previously considered and can occur due to the formation of local secondary structures of RNAs (hairpins), likely to be relatively short and transient compared to the longer stretches of perfectly matching sense-antisense sequences. Hence, the binding of ADAR would be expected to be weaker on these short hairpins, insufficient for multiple editing. Conceivably, only hyper-edited RNA molecules undergo rapid degradation starting from the 3’-UTR (45), thus hampering their quantitative sequencing using the polyA selection strategies, applied herein for the DRS data. This can explain the near absence of hyper-edited DRS reads, except for a cluster limited to a single OsHV-1 genomic region. Although the activity of ADAR during the OsHV-1 infection is now apparent, its biological role remains unresolved. Besides enzymatic activity, we should consider possible editing-independent roles of ADAR, such as the competitive binding to dsRNAs, preventing the action of other RNA binding molecules. Such editing-independent roles can involve miRNA formation (46,47) and RNA degradation, among others (48).

## Conclusions

In this study, we reported a detailed transcriptional map of OsHV-1 based on long Nanopore RNA reads. Undoubtedly, long reads hold a greater potential to provide expression and structural information of overlapping gene architectures, a reason to favour them for (herpes) viral transcriptomics. We also provided the first application of DRS for the detection of inosines in viral RNAs. Although the available tools are not fully developed, they are already promising to untangle inosine modifications. Overall, we showed that the transcriptional architectures of OsHV-1 can likely play a functional role in infection, here highlighted by the detection of the pan-*Herpesvirales* conservation of the capsid scaffold transcriptomic architecture as well as by the presence of “editing decoys”, possibly attracting a considerable part of the enzymatic competence of the host ADAR.

## Materials and Methods

### Experimental infection of *A. broughtonii* with OsHV-1

*A. broughtonii* specimens were collected from a wild population (size range: 56.48 to 68.74 mm) and were cultured for 24 days, fed with homemade shellfish diets, maintained at a temperature of 15.1–15.8 °C, and tested negative to OsHV-1. A viral inoculum was prepared from an infected blood clam collected in a local hatchery, as previously described (12) and 100 μL (adjusted at 10^4^ copies of viral DNA/μL) of the homogenate was injected into the foot of the clams, with paired controls injected with seawater. The infection trial was followed for 72 hours, with physiological conditions monitored every 6-12 hours and DNA, RNA and protein samples collected at 6, 12, 24, 36, 48, 60 and 72 hours post-injection (hpi).

### RNA extraction and high-throughput sequencing

Clam hemolymph was collected from the adductor muscle sinus of three specimens per time point using a 23G needle attached to a 5-ml syringe. The hemolymph samples were centrifuged at 800 rpm for 5 min at 4 °C and, after removing the supernatants, 1 ml of Trizol (Invitrogen, Carlsbad, USA) was added and samples were stored at -80 °C for RNA and DNA extraction. DNA extraction was performed using a TIANamp™ Marine Animals DNA Kit (Tiangen Biotech, Beijing, China) according to the manufacturer’s protocol and used to quantify the OsHV-1 DNA load in the collected samples by quantitative PCR (qPCR), as described previously (49). Total RNA was extracted using Trizol reagent, according to the manufacturer’s protocol and the RNA quality was assessed on an Agilent 2100 Bioanalyzer (Agilent Technologies, Palo Alto, USA).

According to OsHV-1 load measurements, the three biological replicates collected at 60 and 72 hpi were selected for Nanopore direct RNA sequencing. In detail, 800 ng of total RNA was used to produce six libraries (SQK-RNA001, Oxford Nanopore Technologies) according to the manufacturer’s instructions. The poly(T) adapter was ligated to the mRNA using the T4 DNA ligase (New England Biolabs) in the Quick Ligase reaction buffer (New England Biolabs) for 15 min at room temperature. First-strand cDNA was synthesized by SuperScript III Reverse Transcriptase (Thermo Fisher Scientific) using the oligo(dT) adapter and RNA–cDNA hybrids were purified using Agencourt RNAClean XP magnetic beads (Beckman Coulter). The sequencing adapter was ligated to the mRNA using the T4 DNA ligase (New England Biolabs) in the Quick Ligase reaction buffer (New England Biolabs) for 15 min at room temperature, followed by a second purification step using Agencourt beads. Nanopore sequencing of the six libraries was carried out on 6 GridIon flow-cells, to generate approximately 5 million (M) reads per sample.

### Preliminary analysis of DRS reads

Raw electric signals were converted into nucleotides using *guppy v6.2.1* in high confidence mode, merging the resulting fastq files per sample. Critical parameters, such as read length and average quality of the readouts, were evaluated using *Nanoplot v1.40.2*. Nanopore reads were aligned against the OsHV-1 (ID:) and *A. broughtonii* (REF) reference genomes separately, using *minimap2 v2.22* (50) with the following parameters, -ax splice -k14 -uf --secondary=no. The number of mapped reads was counted with *samtools* (51) to evaluate the percentages of on- and off-target reads for each reference.

### Annotation of the OsHV-1 genome with transcripts

The uncorrected DRS reads were initially used to annotate the OsHV-1 genome with transcripts. In detail, *samtools* was used to extract the reads mapped on each OsHV-1 genome strand and separate coverage graphs were generated with *BEDtools* (52). Then, only the 3’ and 5’ mapping boundaries of each read were used to identify transcription start sites (TSS) and transcription termination sites (TTS) using the HOMER *findPeaks* module (53), adjusting - localSize and -size parameters to 50-300 and 20-100 for TSS and TTS, respectively. TSS and TTS occurrences were counted and used to generate a ‘theoretical transcriptome’ annotation file, by combining TSS and TTS found on each strand and populating these intervals with all the possible pseudo-transcripts in a length range of 250-15,000 base pairs (bp). These pseudo-transcripts were used as reference for mapping DRS read. In detail, the DRS reads not mapped on the clam genome were error-corrected using *RATTLE v1.0* (54) and clustered into isoforms. The corrected reads per sample were mapped on the theoretical OsHV-1 pseudo-transcripts with *minimap2* using the following parameters: -ax map-ont -p 0 -N 10. *Samtools* was used to sort and index the alignments in bam format, which were then used as input for *NanoCount* (55). *NanoCount* was run with the -d 50 and -u 50 flags, in order to generate transcript abundance counts considering only reads mapped within 50 bp from TSS and TTS sites. The subset of pseudo-transcripts supported by at least two full-length reads was annotated on the OsHV-1 genome as ‘mRNA’ and used for expression analysis, which was carried out with *NanoCount*, as described above. *Prodigal v2.6.3* (56) was used to predict the ORFs encoded on the validated pseudo-transcripts and generate an OsHV-1 proteome, which was then compared with the available OsHV-1 proteome by reciprocal *blastp* searches. The predicted transcripts were compacted into genes using the *flattenGTF* script (57) and OsHV-1 genome regions with overlapping antisense transcripts were identified with *bedtools intersect* and annotated as ‘dsRNA’. Short-read Illumina datasets were mapped on the OsHV-1 genome annotated with transcripts using the CLC mapped with 0.8 and 0.8 of similarity and length fraction.

### Analysis of ADAR-mediated hyper-editing

The 6 Illumina datasets paired to the Nanopore DRS libraries were used to investigate the extension of ADAR-mediated hyper editing impacting OsHV-1. The *hyperediting* tool (58) was applied after minimal modifications implemented to overcome software incompatibilities of the original version. The parameters were adapted applying: 5 for Minimum of edited sites at Ultra-Edit read (%); 60 for Minimum fraction of edit sites/mismatched sites (%); 25 for Minimum sequence quality for counting editing event (PHRED); 60 for Maximum fraction of same letter in cluster (%); 20 Minimum of cluster length (%); and imposing that the hyper-editing clusters should not be completely included in the first or last 20% of the read. The obtained reads, representing putatively hyper-edited reads, were realigned to the reference genome using the CLC mapper with 0.8 and 0.8 of similarity and length fraction (Qiagen, US). The editing sites were also detected by SNP calling using the SPRINT software (59) using custom parameters. The first 6 bases of each read were trimmed to reduce potential mapping errors. The reads were then aligned with *bwa* v.0.7.12 (60). Paired-end reads were mapped separately with the command options of ‘bwa aln fastqfile’ and ‘bwa samse -n4’, as suggested (61). *Samtools v.1.2* was used for all the format conversions (51), and *picard-tools* v.1.119 was then used to remove PCR duplicates in the sorted BAM files with the command option of ‘MarkDuplicates.jar REMOVE_DUPLICATES = true’ (http://broadinstitute.github.io/picard). The reads with high mapping quality (≥20) were considered as mapped reads and the resulting editing sites where filtered out if possessed less than 5 supporting reads and a frequency lower than 1%. The reads with high mapping quality (≥20) were considered as mapped reads and the resulting editing sites were kept only if possessed both 5 supporting reads and a frequency higher than 1%.

The detection of *inosine-like* modifications in the DRS data was performed with the *DeepEdit* tool (44), with a few modifications described below. The ids of the DRS reads mapped on to the OsHV-1 genome were extracted with samtools from the sam file and used to extract the corresponding data from the fast5 files only using the *ont_fast5_api* (https://github.com/nanoporetech/ont_fast5_api). The fast5 subset was then converted to single fast5 reads files with *multi_to_single_fast5* using standard parameters and used for genome mapping in order to assign the raw electric signals to the final DRS reads using the *tombo re-squiggle* algorithm (62). Finally, the alignment file in sam format was processed using the *DeepEdit* tool (44) in order to obtain a list of validate RNA editing sites with their editing frequencies as well as the positions of inosine-like bases along the DRS reads. A SNV was considered validated if covered by at least four DRS reads gathering the inosine base and showing at least 2 folds more modified reads on the predicted editing strand.

### Proteomic data production and analysis

Proteins were extracted from the samples collected at 6, 24, 48 and 72 hpi. Proteins were extracted using the chloroform-isopropanol method and the final precipitate was re-dissolved with 8 M urea (containing 1% protease inhibitor), and protein concentration was determined by using the BCA kit. Trypsin digestion was performed on an equal amount of each sample protein with a ratio of 1:50 (protease: protein, m/m) overnight. Dithiothreitol (DTT) was added to a final concentration of 5 mM and reduced at 56°C for 30 min, after which iodoacetamide (IAA) was added to a final concentration of 11 mM, and incubated for 15 min at room temperature. HPLC classification was performed by grading the peptides by high pH reversed-phase HPLC on an Agilent 300Extend C18 column (5 μm particle size, 4.6 mm inner diameter, 250 mm length) in a 8%-32% gradient of acetonitrile, pH 9. The peptides were the dissolved in liquid chromatography mobile phase A and then separated using an EASY-nLC 1000 ultra-high performance liquid chromatography (UHPLC) system. The peptide fragments were separated on the UHPLC system and then injected into the NSI Ion Source for ionization and then analyzed by QE mass spectrometry. The ion source voltage was set at 2.2 kV, and the peptide parent ion and its secondary fragments were detected and analysed using a high-resolution Orbitrap. The primary mass spectrometry scanning range was set from 400 to 1500 m/z with a scanning resolution of 70,000.00, while the secondary mass spectrometry scanning range was set at a fixed starting point of 100 m/z and the secondary scanning resolution was set at 17,500.00. The data acquisition mode used a data-dependent scanning (DDA) procedure. Secondary mass spectrometry data were searched using Maxquant 1.5.2.8. The search parameters were set as follows: the database was composed by the blood clam and OsHV-1 proteomes (24172+ 132 sequences), an inverse library was added to calculate the false-positive rate (FDR) caused by random matches, and common contaminant libraries were added to the database for eliminating the influence of contaminant proteins in the identification results; the digestion method was set as Trypsin/P; the number of missed cut sites was set as 2; the minimum length of peptide was set as 7 amino acid residues; the maximum number of peptide modifications was set to 5; the mass error tolerance of primary parent ions for First search and Main search was set to 10.0 ppm and 5 ppm, respectively, and that of secondary fragment ions was set to 0.02 Da. Carbamidomethyl(C), a cysteine alkylation, was set as a fixed modification. The quantification method was set to TMT-6plex, and the FDR for protein identification and PSM identification were set to 1%. A total of 798,016 secondary spectra were obtained by mass spectrometry analysis. After searching the protein theoretical data, the number of available effective spectra was 140,218, with a utilisation rate of 17.6%. A total of 72,876 peptides were identified by spectral analysis, including 71,478 specific peptides originating from 5,083 Arc clam and 41 OsHV-1 proteins.

### Database searches and structural modelling

For the annotation of OsHV-1 proteins, similarity searches in databases has been performed as follows. Blastp and iterative blastp searches (PSI-BLAST) against the NCBI nr clustered database were performed using 10E-10 as cutoff E-value to consider the hits for the subsequent iterations. To detect distant homology, the *HHblits* v3.3.0 tool (63) was used in combination with the pdb70 database. Functional annotations have been retrieved using *InterProScan* v.5.57-90.0 (64).

### Data rendering and statistical analysis

OsHV-1 genomic features, annotations and coverage levels were inspected and rendered using the GVIZ package (65) implemented in *R v4.2.1* (66). All the plots and statistical analyses have been performed using the *tidyverse*, *ggplot2*, *ggpubr* and *smplot2* packages and with CLC Genomics Workbench (Qiagen, US).

## Acknowledgments

Part of the computational power required for the analyses was provided by “University of Padova Strategic Research Infrastructure Grant 2017: “CAPRI: Calcolo ad Alte Prestazioni per la Ricerca e l’Innovazione”.

## Supporting information captions

Table S1. Annotation of the OsHV-1 proteins. For the 132 OsHV-1 proteins the ID, the length in number of residues, the functional annotations obtained using *Interproscan* (database, annotation ID, description, positions along the protein, Interpro ID, description and associated GO terms) and the *blastp* best match (accession ID, the E-value, the description) are reported.

Table S2. Expression levels of OsHV-1 transcripts in the 60 and 72 hpi samples detected by short-(ILL) and long-read (ONT) technologies. Expression levels are indicated as Transcripts Per Million (TPM).

Table S3. List of the 465 ADAR-compatible SNVs tested for the presence of inosines. The table indicated the SNV position, the reference and allele base, the eventual occurrence within coding regions and the amino-acid substitution, the presence of other variants in the same codon and if SNVs are synonymous or non-synonymous as well as if they are validated for the presence of inosine.

Figure S1. Percentages of OsHV-1 reads over total reads measured along the infection time-course experiment (three replicates per time point, reported as triangles in the box plot).

Figure S2. Distribution of the transcript lengths (measured in base pairs) calculated for 65 gene units, normalized by the gene unit length. The gene units with at least 100 mapped DRS reads were selected.

Figure S3. Distributions of the transcript expression levels (Transcript Per Million, TPMs) measured using DRS data and Illumina data in the six samples. Correlation R and the associated p-value are reported on the graphs.

Figure S4. Heat map reporting the expression levels of the transcripts (log10 scale) measures using Illumina data along the time-course infection experiment (36-72 hpi). Above the heat map the clustering of the samples is reported and their identity is color-coded.

Figure S5. The box plot depicted the percentage of hyper-edited reads over total rads measured at 60 and 72 hpi (a) and the percentages of DRS reads with at least one detected inosine nucleotide at the same time points (b).

## Data availability

The data described in this work are available in public repositories or are provided within supplementary materials. In details, long read and short read data are available in the NCBI SRA archive under accession IDs ‘PRJNA874858’. Proteomic data are deposited in the iProX (https://www.iprox.cn/) archive, under project ID IPX0007584002.

## Author’s contribution

UR: conceptualization, data analysis, writing, discussion, review and editing. EB: data analysis. BWH: methodology. XZ: methodology. LSX: validation, resources. MK: conceptualization, data analysis, writing, discussion, review and editing. CMB: funding acquisition, conceptualization, writing, review and editing.

